# Reinforcement and selection against migrants maintain sperm competitive and genetic differences among populations of *Drosophila pseudoobscura*

**DOI:** 10.64898/2025.12.29.696855

**Authors:** Dean M Castillo, Leonie C. Moyle

## Abstract

Sexual interactions among closely related species can select for divergence in reproductive traits as a mechanism to avoid the costs of hybridization. This selection, termed ‘reinforcement’, produces a canonical pattern of stronger isolation in populations that are sympatric with congeners, versus in allopatric populations. While reinforcement has been observed in many systems, the resulting pattern might be transient. There must be selection against migrants between allopatry and sympatry—akin to local reproductive adaptation—for a pattern of stronger sympatric isolation to be maintained. We tested for evidence of this selection against migrants in the context of sperm competition, in *Drosophila pseudoobscura*. Previously we showed an advantage of sympatric genotypes in sympatric contexts, via their elevated reproductive isolation (conspecific sperm precedence) against congeneric *D. persimilis*. Here, using experimental crosses that simulated migration between populations that are naturally sympatric and allopatric with *D. persimilis,* we show that sympatric genotypes are at a sperm-competitive disadvantage against allopatric male genotypes. This selection against sympatric males in allopatry, combined with the advantage of sympatric genotypes in sympatry, produce a scenario for strong selection against migrants between these two geographic contexts. This is accompanied by population differentiation in reproductive gene expression and allele frequency variation between allopatry and sympatry, including genes with known roles in sperm competition. While some of these changes are parallel among sympatric or allopatric contexts, most are unique to individual populations and might reflect the unique ways that selection has acted on sperm competition genes in different sympatric populations.

## Introduction

Because sexual interactions are shaped by the identity of local mating partners, spatial variation in the makeup of local mating pools can result in differentiation among populations (Lackey et al., 2024; Maan & Seehausen, 2011; Tinghitella et al., 2013). This differentiation in sexual interactions is maintained when selection on mating interactions is stronger than the ameliorating forces of migration and gene flow (Servedio & Noor, 2003; Yukilevich & True, 2006). One such context for strong selection on local mating interactions is when species co-occur (i.e. are sympatric) with a closely related sister species. Under these conditions, selection can act to increase the frequency of variants that reduce costly heterospecific matings—the phenomenon referred to as reinforcement (Butlin, 1987; Coyne & Orr, 2004; Howard, 1993). Reinforcement produces a specific pattern of trait divergence between populations, with sympatric populations exhibiting stronger reproductive isolation compared to allopatric populations (Noor, 1999; Servedio & Noor, 2003). Additionally, because co-occurrence with heterospecifics in sympatric populations alters the local composition of potential mates it can also can shape within-population sexual interactions (Castillo & Moyle, 2019; Hoskin et al., 2005; Silvertown et al., 2005). Reinforcement is therefore a powerful context in which to understand the contribution of strong local selection to shaping sexual differentiation between conspecific populations. However, while many studies of reinforcement focus on the pattern of trait differentiation between sympatric and allopatric populations, we do not have compelling examples of the factors that maintain these differences in the face of homogenizing gene flow between these two geographical contexts.

In the presence of migration and gene flow, differentiation in reproductive traits can be maintained between sympatric and allopatric populations if selection against migrants is strong, analogous to local adaptation where migrants are reproductively maladapted (Anderson et al., 2023; Comeault & Matute, 2016; Yukilevich & Aoki, 2016). Selection against allopatric alleles in sympatric populations can come from reinforcement itself, as allopatric populations lack a history of strong selection for variants that reduce heterospecific hybridization. However if there is no selection against sympatric migrants in allopatric populations, sympatric alleles should increase in allopatric populations over time (Comeault & Matute, 2016; Yukilevich & Aoki, 2016). This export of sympatric alleles would thereby erase the allopatric/sympatric contrast that is the discernable pattern of reinforcement (Comeault & Matute, 2016; Yukilevich & Aoki, 2016). To observe sustained differences between sympatric and allopatric populations, sympatric alleles must have a cost in allopatric populations (Bolnick & Nosil, 2007; Hendry, 2004). Systems where we currently observe reinforcement are therefore powerful models to understand how selection can maintain conspecific population differentiation in reproductive traits.

Here we focus on a case of reinforcement involving changes in sperm competition. Sperm competition manifests as reproductive isolation when conspecific male sperm outcompete the sperm of heterospecific males and sire the majority of the progeny—known as conspecific sperm precedence (CSP) (Howard, 1999; Manier et al., 2013; Yeates et al., 2013). CSP is a potentially strong interspecific barrier because it reduces the formation of hybrids (Castillo & Moyle, 2019; Howard, 1999; Turissini et al., 2018), and has recently been identified as a target of reinforcement (Castillo & Moyle, 2019; Matute, 2010). Moreover, CSP shares a genetic basis with intraspecies sperm competition in *D. melanogaster* (Castillo & Moyle, 2014). As a result, reinforcement of CSP could have potential consequences for sexual interactions within a species, if variants responding to reinforcing selection have pleiotropic effects on competitive interactions with conspecifics. Finally, sperm competition and gamete interactions are known to experience strong selection within populations, including co-evolutionary interactions where male ejaculate and proteins interact with female reproductive tract (Wigby et al., 2020). Therefore sperm competition is one reproductive trait for which reinforcing selection could potentially have direct or indirect effects on the origin and maintenance of conspecific population differentiation in reproductive traits.

Examining sperm competition in the context of reinforcement can also be used to understand the genetic basis of this response, including whether it has drawn on shared or different genetic variants across sympatric populations. The genetic basis of sperm competition variation is understudied in natural populations, however individual genes can have large effects on sperm competition outcomes under experimental conditions (Civetta & Ranz, 2019; Hopkins & Perry, 2022), including members of the sex peptide network (Findlay et al., 2014; McGeary & Findlay, 2020). The genetic complexity of sperm competition can in turn have consequences for whether local sperm competition responses, including CSP, draw upon the same genetic variants in response to local reproductive partners. Studies examining reinforcement often treat sympatric populations as homogenous in terms of the genetic composition of responding sympatric populations and the selection pressure imposed by heterospecific interactions (Fitzpatrick et al., 2008; Foote, 2018; Mallet et al., 2009). However, ecological studies of other species interactions repeatedly show that they can vary across sympatric ranges (Aldridge, 2005; Larson et al., 2013; Travis, 1996; Zeng et al., 2011). Similarly, while a common selection pressure can result in parallel phenotypic and genetic changes when selection acts on standing variation (Bohutínská et al., 2021; Fang et al., 202), parallel phenotypic changes need not be underpinned by parallel genetic changes (Westram et al., 2014). Therefore while we expect that reproductive genes involved in CSP should be differentiated between sympatric and allopatric populations, it is unclear whether these variants would be the same in different sympatric populations.

In this study we used populations of *Drosophila pseudoobscura* that exhibit reinforcement acting on CSP to examine evidence for mechanisms maintaining sustained differences between allopatric and sympatric contexts. The *D. pseudoobscura* system was initially described as a model for reinforcement focused on premating isolation and female choice (Noor, 1995) but more recent observations demonstrate reinforcement now acts on CSP in *D. pseudoobscura*(Castillo & Moyle, 2019). Here we use this system to examine: 1) whether males from sympatric populations suffer from a competitive disadvantage against allopatric males, 2) if this pattern of competitive deficit is consistent across two different sympatric populations, and 3) the extent of genetic differences (gene expression and sequence variation) in reproductive genes between sympatric and allopatric populations. Using sperm competition experiments, we simulated migrant males competing against local males in both sympatric and allopatric populations. We demonstrated that males from sympatric populations were at a sperm competition disadvantage when competing against males from allopatric populations, but not when competing against other sympatric population males. This phenotypic divergence in sperm competition was accompanied by differentiation between sympatric and allopatric populations in expression and allele frequency variation at reproductive genes. Sympatric populations had both parallel and unique changes in gene expression and allele frequency differentiation compared to allopatric populations, consistent with divergent selection in these populations. The loci underlying this pattern of differentiation between populations were enriched for reproductive genes. Overall these data provide evidence for a cost of sympatric alleles in allopatric populations that could produce stable patterns of reinforcement for CSP and maintain population differentiation in reproductive loci, including those involved in CSP and sperm competition.

## Methods

### Wild type fly stocks

The *D. pseudoobscura* stocks used in this experiment were previously described in Castillo and Moyle (Castillo & Moyle, 2019) and have been deposited with the National Drosophila Species Stock Center (Cornell University; Supplemental Table 1). These stocks were initially collected and established as isofemale lines in the summers of 2013 and 2014 from four populations. Two of the populations— near Zion National Park, UT and Lamoille Canyon, NV—are considered allopatric with respect to the closely related species *D. persimilis*. The other two populations—Mt. St. Helena, CA and ‘Sierra’ (near Meadow Vista and Forest Hill), CA—are sympatric with respect to *D. persimilis* and co-occur with this species. All stocks were reared on standard media prepared by the Bloomington Drosophila Stock Center, and were kept at room temperature (∼22C).

### Interpopulation sperm competition assay

To determine the success of males as migrant sperm competitors across sympatric and allopatric contexts, we completed pairwise crosses between males from all four populations. These crosses were conducted in the fall and summer of 2015. Sperm competition assays require that an individual female mate sequentially with two distinct male genotypes. For these experiments, for each individual replicate we used one male that came from the same population as the female (the “local” male) in combination with a male that came from one of the three other populations (the “migrant” male). Each migrant male carried a dominant green fluorescent protein (GFP) marker (described in Castillo and Moyle 2019) to enable quantification of competitive siring success via offspring phenotyping. In all experimental crosses, females were paired first with the local male and subsequently with the migrant male. This assay evaluates the offensive sperm competitive ability of migrant males to displace local male sperm (second male siring ability or P2; (Clark et al., 1995; Gromko et al., 1984). We focused on offensive sperm competition so that we could compare these between population results to within population results (previously published in (Castillo & Moyle, 2019)) which used the same scheme and were completed at the same time as the interpopulation crosses. Based on within population observations, we expect that the second male will sire an average 75-80% of the progeny (Castillo & Moyle, 2019) so we are able to assess evidence for both increases or decreases from this average value in our interpopulation experiment.

For each population-by-population comparison we included four female genotypes from the focal population, four local male genotypes, and two migrant male “tester genotypes”. The female and local male genotypes were represented in a 4×4 design and within these 16 crosses we attempted to use each migrant male tester genotype equally. We ensured that each female x local male x migrant male combination was present in each block of the experiment. Each block contained 128 replicates and the total number of replicates was 512 across all blocks. The intrapopulation crosses had n=64 for each population; across four populations this resulted in 256 crosses. The remaining 256 of 512 were divided between the interpopulation combinations, with the ‘direction’ of each combination determining which population was considered migrant versus focal. Crosses between the two allopatric populations had n=64 (n=32 in each direction) and crosses between the two sympatric populations had n=64 (n=32 in each direction). Crosses between each allopatric and sympatric population combination had n=32 (n=16 in each direction). There are four allopatric-sympatric combinations yielding n=128 total crosses between allopatric and sympatric populations.

For each experimental block, virgin individuals were collected and kept in same sex vials 7 days prior to the initiation of crosses. One day before mating, local males were isolated individually (Castillo & Moyle, 2019; Dixon et al., 2003). The following day, females were individually added (without anesthesia) to a vial containing a local male and were co-housed for 24 hours, after which time the male was removed. We kept females housed individually in these vials for 7 days before the second mating. After 7 days we inspected all vials for the presence of larvae to determine if females had mated with the first male. Only females that had mated (i.e. had produced larvae within 7 days) were retained for the remainder of the experiment. It is unlikely that females remated, or mated more than once, within this first 24 co-habitation period as the average remating rate for *D. pseudoobscura* is 4 days (Snook & Markow, 2001), and we previously found that for these genotypes most would not remate until closer to 7 days (Castillo & Moyle, 2019; Peckenpaugh & Moyle, 2024)

For the second mating, each individual female was paired with one of the migrant male genotypes. These second males were also isolated one day before the introduction of the female. Seven days after mating with the first male, females were transferred, without anesthesia, to the vial containing the second male. Individual pairs were co-housed for 24 hours and the male was removed on the second morning. The female was kept for five days (transferring after 2 days to avoid overcrowding of larvae). All progeny produced in the five-day window after the second mating were collected and scored for the presence or absence of the GFP marker. Our measure of sperm competition (P2) was then the number of wild-type (non-GFP) progeny out of the total number of progeny scored for a particular cross. If all progeny produced in a cross were wild-type (non-GFP), we did not use this replicate because we could not ensure that a second mating had taken place. Individual progeny were scored as they eclosed, using a Leica M205FA Stereo Microscope that has an Hg fluorescent lamp attached and GFP filter. Individuals were anesthetized and the ocelli were examined for GFP signal as described in Castillo and Moyle (Castillo & Moyle, 2019).

### Intrapopulation sperm competition assay

The same assay described above had also been used to estimate intrapopulation sperm competition. While these data have been previously published (Castillo & Moyle, 2019) the crosses were completed at the same time and in parallel as the interpopulation assays described here. The intrapopulation experiments involved a local male competing against a male from its own population that carried the GFP marker. For the purpose of the present study, these assays captured local vs local competition.

### Statistical analysis of sperm competition

In this analysis we wanted to determine if the average sperm competition for migrant males competing against local males was different than local males competing against each other. The local vs local competition data from Castillo and Moyle (Castillo & Moyle, 2019) was used as a baseline for our comparisons. We tested for changes in sperm competition using binomial regression since the proportion of GFP to wild-type progeny is naturally a binomial variable. We ran a separate model for each local population and included the identity of the local male (first to mate) and the female genotype as random effects. In previous studies we have used this model framework to capture the effects of these genotypes on the outcome of sperm competition, (Castillo & Moyle, 2019). The model is of the form

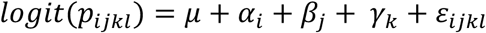

The variable μ is the baseline sperm competition of the local male competing against another local male (see above). For each model the local male was used as the baseline to compare against the other populations. The variable α is a categorical variable with three levels representing the three potential migrant populations. The variables β and γ are also categorical variables with four levels each that represent the local second male genotypes and female genotypes respectively.

The total number of replicates for each population type (intrapopulation vs interpopulation) was different (see above). To compare our estimates across these cross types we needed to take into account the differences in sample size because these differences could lead to differences in variance between cross types and influence our inference (Fithian & Hastie, 2014). Moreover, unequal sample sizes can bias estimates in groups with the smallest sample size, increasing type I errors (Rusticus & Lovato, 2014). Our smallest groups were the migrant allopatric-sympatric crosses, which were of particular interest. Thus, to avoid potential increases in type I errors we performed subsampling of the data to ensure equal sample size across all cross types (Politis et al., 2001). Combinations that had larger sample sizes were subsampled with replacement to the smallest number of replicates (n=16). Combinations that already had n=16 (i.e. allopatric-sympatric combinations) were also sampled with replacement. To capture all of the biological variation that existed in each cross type, we subsampled the entire dataset 1000 times and ran each model on these subsampled datasets. A significant difference from the baseline was determined by comparing the 95th percentile confidence intervals for the coefficients, with these intervals generated from the distribution of coefficients that were fit from each subsampled replicate (Politis et al., 2001). Subsampling has the potential to decrease power to the smallest sample size, thus our resulting estimates of differences among cross types are conservative.

### Genetic differences between populations and divergence from D. persimilis

To generate data on both sequence variation (single nucleotide polymorphisms - SNPs) and variation in gene expression across these populations of *D. pseudoobscura*, we sequenced whole body transcriptomes for all wild-type strains used in this experiment. We also sequenced wild-type sympatric *D. persimilis* and two additional wild-type strains of *D. pseudoobscura* per population. These extra *D. pseudoobscura* were included so that our total dataset included six *D. pseudoobscura* strains per population. Our goals with these data were to 1) quantify the genetic differences between sympatric and allopatric populations and 2) determine which of these changes corresponded to differences in reproductive genes. In addition we used these data to quantify enrichment of sex biased genes, test for selection on differentiated loci and quantify which changes were parallel and shared between the sympatric populations.

To generate these data, transcriptomes were prepared by extracting total RNA from 20 whole bodies from each sex within each strain, to generate separate sex-specific RNAseq libraries for each strain. There were six male strain and six female strain samples for each of the four *D. pseudoobscura* populations, and four male strain and four female strain *D. persimilis* samples, resulting in 56 libraries total. To generate these transcriptomes, virgin individuals were collected and maintained in same sex vials for seven days. Twenty flies were placed, without anesthesia, into an Eppendorf tube and immediately submerged in liquid nitrogen and stored at - 80C. For extraction, tubes were removed from −80C, placed on ice, and samples were immediately homogenized with Trizol reagent. Total RNA was extracted using the Invitrogen PureLink RNA mini kit (ThermoFisher Scientific). Sequencing libraries were prepared by the IU Center for Genomics and Bioinformatics (CGB). Libraries were sequenced by the CGB using two runs of 75bp long reads on the Illumina NextSeq (Illumina). This generated ∼3.5million reads per sample. Raw reads were processed by removing adapters using scythe (Buffalo, 2011) and quality trimming using sickle (Joshi & Fass, 2011). Reads were mapped to the *D. pseduoobscura* r3.04 genome using bwa mem (Li, 2013). Mapping reads from *D. persimilis* to the *D. pseudoobscura* genome did not affect downstream analysis as ∼98% of the reads mapped across samples, similar to the mapping of *D. pseudoobscura.* PCR duplicate reads were identified using picard (Broad Institute) and filtered for quality score above 20 using samtools 1.8 (Li et al., 2009).

From these filtered and mapped reads, counts for each gene were determined using featureCounts (Liao et al., 2014)for each library separately (n=56). Therefore, gene expression variation and differentiation is treated separately for males and females in each population. Before calling SNPs, male and female reads from the same strain were merged using samtools (therefore sequence variation and differentiation is analyzed without regard to sex). SNPs were then called using a best practices GATK pipeline that included the haplotype caller algorithm followed with joint calling protocol (DePristo et al., 2011). All raw reads are available through the SRA with accession numbers provided in Supplemental Table 1.

### Expression differences and divergence between sympatric and allopatric populations

We tested whether there were differences in gene expression across populations by comparing gene expression levels in sympatric populations to two allopatric populations, and comparing all *D. pseudoobscura* populations to *D. persimilis.* We tested for differences in expression between *D. pseudoobscura* populations and between *D. pseudoobcura* and *D. persimilis* following the workflow used in Fuller et al (Fuller et al., 2016). We used EDAseq to correct for GC content bias in the transcriptome (Risso et al., 2011)and accounted for other sources of technical (i.e. not biological) variation in the dataset using RUVSeq (Risso et al., 2014). Finally, upper-quartile normalization was performed on the filtered read counts using the R package RUVSeq (Risso et al., 2014). Normalized read counts were used in models with edgeR that incorporates the matrix containing normalization offsets and the matrix from RUVseq (Risso et al., 2014).

We used a principal components analysis to explore large scale differences and clustering between species and populations (Fig 1), and followed this with specific comparisons to test for expression differences between species and between sympatric and allopatric populations. We first quantified expression differences between each *D. pseudoobscura* population and *D. persimilis*. This allowed us to determine if there were expression differences distinctive to sympatry, compared to allopatry, when contrasted with its sister species. We then compared each sympatric population individually to the allopatric populations. This allowed us to quantify expression differences that occurred in that specific sympatric population.

**Figure 1.**
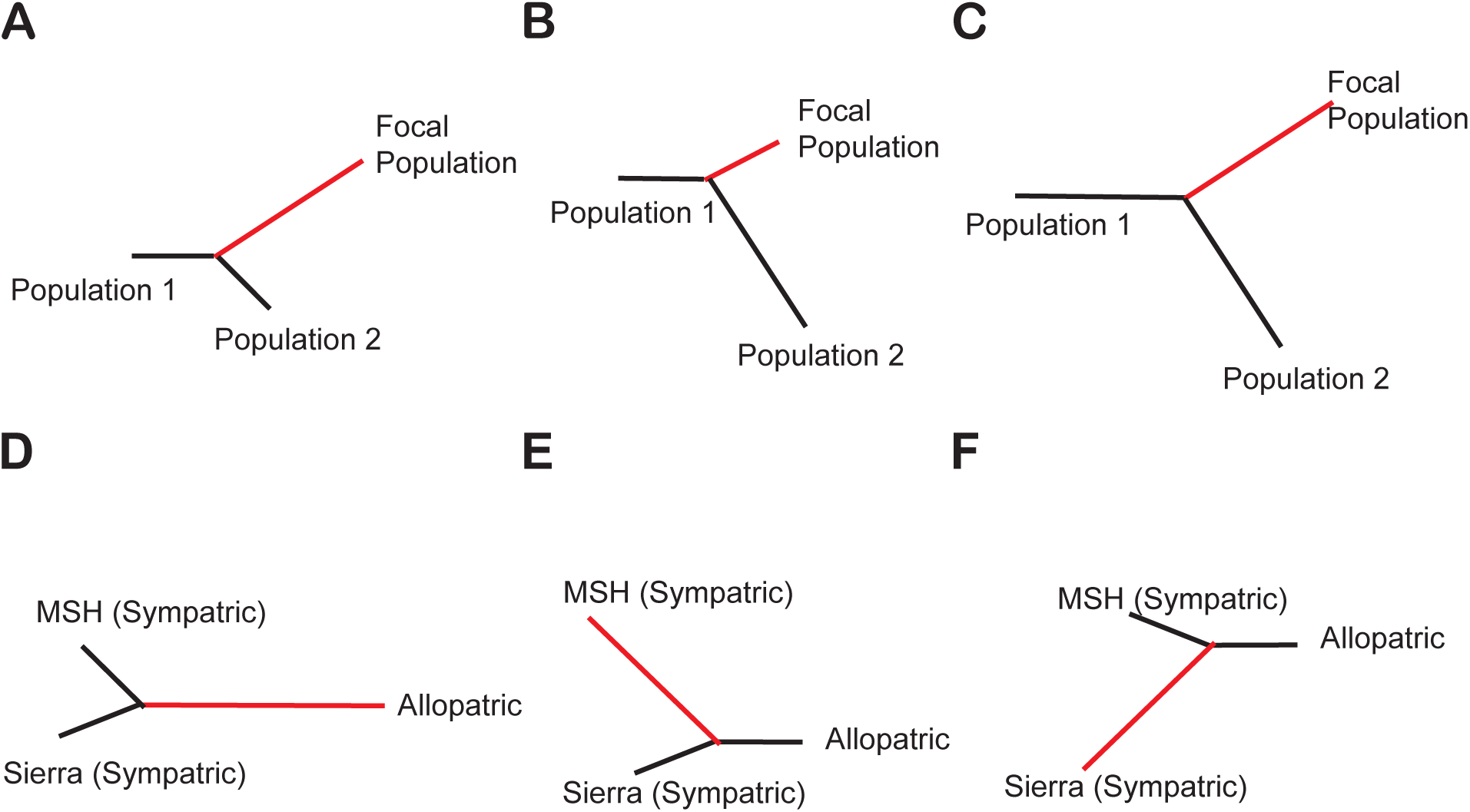
Panels A-C show hypothetical trees with 3 populations. Panels D-F show the actual tests used in this study. A) The focal population has experienced evolution at a specific locus, as evidenced by increased Fst compared to the other two populations; the resulting longer branch produces a PBE outlier. B) The focal population has not experienced selection at this locus, and would not be a PBE outlier. C) Evolution/selection has occurred in the focal population, but also in the other two populations. This scenario would not be a PBE outlier. D) The focal population is the allopatric population allowing us to identify selection that differentiates sympatric from allopatric populations. E) MSH is the focal population. This allows us to identify selection unique to this sympatric population. F) Sierra is the focal population. This allows us to identify selection unique to this sympatric population. In cases E and F different sites within the same gene could be under selection creating selection on shared loci.

Finally, we tested whether the differences (from allopatry) in expression observed in each of the two sympatric populations were parallel and occurred more often than would be expected by chance using a proportion test (Jacobs et al., 2020; Rivas et al., 2018). We compared the observed number of genes that were differentially expressed in the same direction in both sympatric populations compared to allopatric populations, to the expected number of differentially expressed genes from a randomization test. In the randomization test, samples were drawn based on the significant number of genes in each comparison and the expected value was the mean number of shared genes across samples (Jacobs et al., 2020; Rivas et al., 2018).

### Sequence variation and differentiation between populations and species

To identify SNPs and genomic regions that showed allele frequency differences, we used vcfTools (Danecek et al., 2011)to calculate pairwise Fst between all populations for each SNP and a genome wide average Fst. Using these pairwise Fst estimates, we calculated the Population Branch Excess (*PBE*) statistic (Pool et al., 2017; Yassin et al., 2016). to identify variants consistent with local selection. The benefit of the *PBE* approach, compared to other outlier methods, is that it allows us to estimate differences in allele frequency due to selection in specific populations, where a “population” can be any grouping of individuals of interest (Coughlan et al., 2021; Wang et al., 2021). To do so, the *PBE* statistic uses three branches, assigning one branch as the focal branch. The statistic is conditioned on the changes between the non-focal populations, thereby accounting for any evolution occurring between the two non-focal populations and providing directionality to the statistic (Pool et al., 2017; Yassin et al., 2016)). We focused our analysis on individual SNPs rather than windows, that would average *PBE* across sites, because *PBE* has increased performance for identifying individual SNPs potentially underlying local adaptation (Coughlan et al., 2021; da Silva Ribeiro et al., 2022; Shpak et al., 2024).

We used three tests as described below (Fig 1), to determine genetic differences between sympatry and allopatry, and whether changes in sympatry were shared. Since the *PBE* approach is limited to three branches, for each test we combined both allopatric populations into a single population and kept the two sympatric populations separate. Therefore, in all three of our tests the three groupings (“populations”) are constant, while the focal population differs (Fig 1). In the first test, we made the combined allopatric population the focal population; this allowed us to quantify differences that were shared in sympatric populations by isolating the branch between the allopatric vs sympatric populations (Fig 1D). In the other two tests we alternated which sympatric population was the focal population. This allowed us to quantify the changes that were unique to each sympatric population (Fig 1E-F). To examine shared PBE loci we used the method described above for DE genes (Jacobs et al., 2020; Rivas et al., 2018).

After calculating the *PBE* statistic for each SNP we retained the top 1% of outliers and tested for evidence of enrichment of reproductive genes. We used three distinct approaches to identify reproductive genes for these analyses, that leveraged three different sources of available reproductive data. In the first we used a gene ontology (GO) analysis based on *D. melanogaster* orthologs. In the second we calculated tissue biased patterns of expression from previously-published tissue-specific transcriptome data from *D. pseudoobscura* and used these categories to test for enrichment. In the last approach we used loci previously identified as being present in the seminal proteome of *D. pseudoobscura* as our set of genes to test for enrichment.

For the first analysis, we created a list of PBE loci that had orthologs with *D. melanogaster*. We used Pangea to test for enrichment using the SLIM GO terms and Flybase mutant annotations (Hu et al., 2023). For the second analysis, we used adult tissue specific transcriptome data from Yang et al (Yang et al., 2018) to calculate the statistic tau (Yania et al., 2005)for all genes in the *D. pseudoobscura* genome. This statistic ranges from 0 to 1, with tau=1 indicating a gene is expressed exclusively in a single tissue. The transcriptome data came from 12 tissues total. Eight of these were non reproductive tissues and included male and female abdominal carcass, male and female thorax, male and female head, and male and female digestive tract. The four reproductive tissues were male testes, male accessory gland and ejaculatory bub, female ovaries, and female reproductive tract (bursa, spermathecae, seminal receptacle). We used a cutoff of tau=0.75 to assign biased expression to a specific tissue. Thus a gene was considered reproductive if it had a tau of 0.75 with highest expression in one of the four reproductive tissues. In our third approach, we used a list of loci found to be present in the male accessory gland and transferred to the female reproductive tract, as determined by high resolution proteomics (Karr et al., 2019). We categorized our PBE loci as being present vs absent in the proteomics list and compared our PBE loci to the total number of loci in the list to assess for enrichment. Note that the proteomic list only partially overlapped with reproductive genes identified using the tau statistic, which could reflect both biological and technical differences. For this reason we chose to look for enrichment in both categories separately.

Lastly, we tested if any male reproductive genes have experienced longer-term selection that could contribute to reproductive differences (i.e. CSP) between species. We focused on male reproductive genes because we have the most comprehensive data—from both tissue specific expression and proteomics—for their identification. For these tests, we used the transcriptomic data to generate coding sequence alignments for loci were both PBE outliers and were also considered a male reproductive gene. Using these alignments we determined the number of segregating and fixed differences between *D. pseudoobscura* and *D. persimilis* and calculated McDonald Kreitman statistics (McDonald & Kreitman, 1991). This allowed us to identify genes that could respond to selection in sympatry and contribute to species differences.

### Patterns of differentiation in known seminal fluid proteins

In addition to our genome-wide PBE outlier analyses, we also evaluated patterns of allelic variation in a set of *a priori* candidate loci known to be involved in sperm competition, since they might contribute to sperm competition and CSP (Castillo and Moyle 2014; Castillo and Moyle 2019.), including in these populations. To do this we evaluated allelic frequency variation in all genes that have orthologs in the sex peptide network (Findlay et al., 2014; McGeary & Findlay, 2020). While some of these genes contained SNPs that were PBE outlier loci, most did not. This is not unexpected given how sperm competition genes often maintain variation that is different from other loci under selection between populations (Clark, 2002; Kern et al., 2004) as well as how the PBE test is constructed (Pool et al., 2017; Shpak et al., 2024). If Fst is high between multiple branches of a gene tree simultaneously, the PBE statistic would not identify these loci as outliers (Pool et al., 2017; Shpak et al., 2024), because the signal would not be specific to a focal branch (Fig 1D). To overcome this, we used pairwise Fst estimates focusing on SNPs within sex peptide network genes and comparing them to the genome wide threshold for each specific pairwise comparison. This allowed us to identify cases where selection could be acting in multiple populations, and would not show up in the PBE outlier test. A similar approach using Fst has been used to identify signals of recent selection on reproductive proteins in humans (Schaschl & Wallner, 2020).

We asked if there were SNPs within each locus that was in the 99% percentile of Fst for that population comparison. Outliers were determined using the specific Fst distribution for each pairwise combination. To test if the Fst observed in our sperm competition loci was greater than expected by chance, we generated 1000 random samples. We chose 28 loci randomly and retained the SNP with the max Fst per population comparison, repeating this 1000 times. We compared the distribution of these Fst values to the mean Fst of the sperm competition loci for each comparison to generate an empirical p-value.

## Results

### Sympatric migrants are poor competitors in interpopulation sperm competition

We compared sperm competition between local males and migrant males from each population. Although there was variation between sympatric populations, in general we found that migrant males from sympatric populations were less competitive against local allopatric males compared to the allopatric intrapopulation crosses (Fig 2; Table 1). Specifically when the allopatric Lamoille population was the focal population, migrant males from sympatric Sierra were significantly worse at sperm competition compared to the intrapopulation baseline (*β* = (−1.0950, −0.1210), 95% confidence interval). When the allopatric Zion population was the focal population, migrant males from both sympatric Mt St Helena and sympatric Sierra populations were significantly worse sperm competitors (MSH *β* = (−1.4676, −0.1313), Sierra *β* =(−1.5891, - 0.2222)) (Fig. 2; Table 1). This poor sperm competitive ability as migrants is also reflected in sympatric populations’ lower baseline (intrapopulation) P2 against their own males. Since this lower baseline P2 is observed in both sympatric populations, they do not have a significant ‘migrant’ effect in the context of each other (Fig 2). In contrast, migrant males from allopatric populations into sympatric contexts trend towards having a competitive advantage against local sympatric males, although this was not statistically distinguishable from the local baseline proportion in both sympatric contexts (Fig 2; Table 1). This is consistent with the observation that males from both allopatric populations have a higher baseline P2 within their own population.

**Figure 2.**
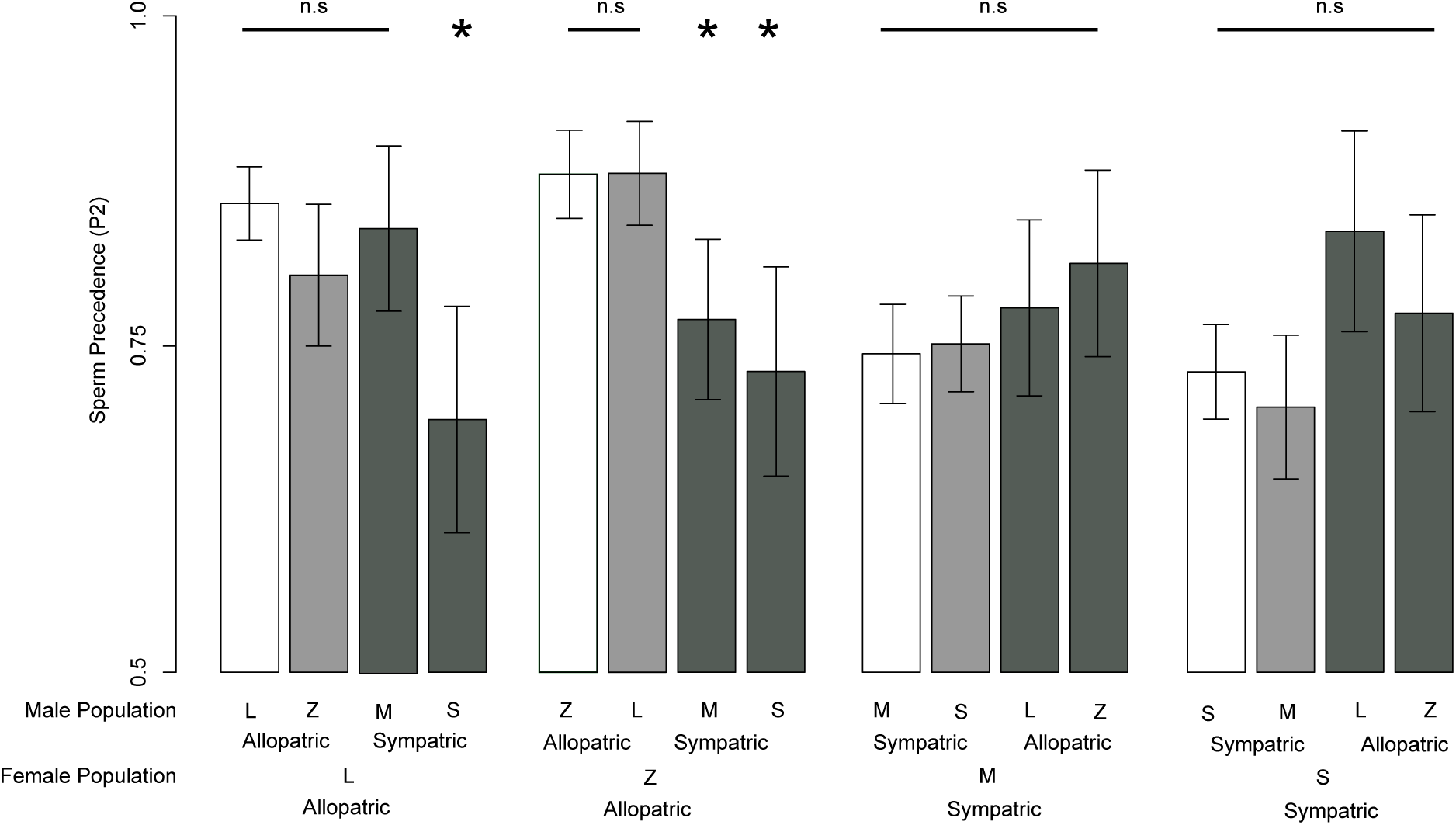
Sperm precedence, the percent of progeny sired by the second male, for migrant males across populations. For each female population the male from the same population is on the left (white) and is considered baseline in the statistical model. The next bar is a male from a different population but similar geography, allopatric or sympatric, (light grey) followed by two populations with males from the opposite geography (dark grey). * indicates non-overlapping confidence intervals in sperm precedence compared to the baseline for each model.

**Table 1.**
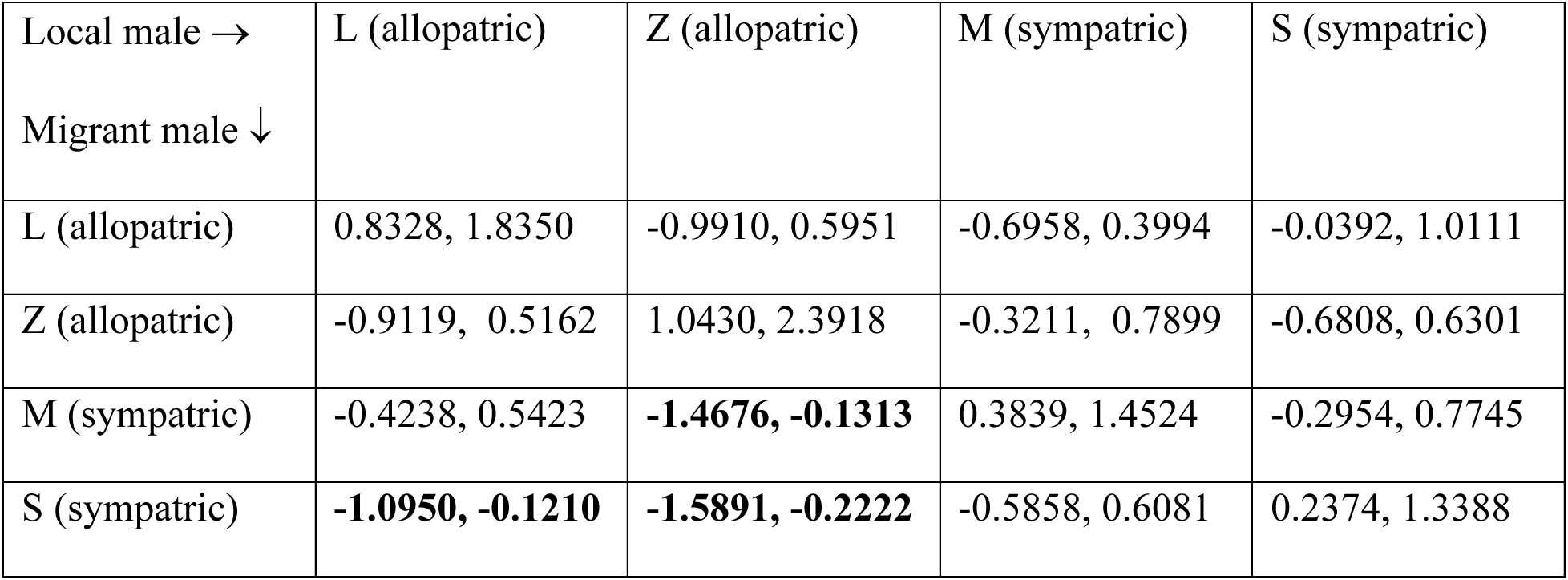
Sympatric males suffer in sperm competition when they are the migrant male. Confidence intervals for sperm competition (P2) for migrant males compared to the within population estimate (diagonal). Confidence intervals that do not overlap zero for migrant male population level effects are in bold. Within population estimates (diagonal) have confidence intervals uniformly greater than zero, consistent with standard second male (P2) precedence.

| Local male →<br>Migrant male ↓ | L (allopatric) | Z (allopatric) | M (sympatric) | S (sympatric) |
| --- | --- | --- | --- | --- |
| L (allopatric) | 0.8328, 1.8350 | -0.9910, 0.5951 | -0.6958, 0.3994 | -0.0392, 1.0111 |
| Z (allopatric) | -0.9119, 0.5162 | 1.0430, 2.3918 | -0.3211, 0.7899 | -0.6808, 0.6301 |
| M (sympatric) | -0.4238, 0.5423 | **-1.4676, -0.1313** | 0.3839, 1.4524 | -0.2954, 0.7745 |
| S (sympatric) | **-1.0950, -0.1210** | **-1.5891, -0.2222** | -0.5858, 0.6081 | 0.2374, 1.3388 |

### Sympatric populations primarily have unique differences in gene expression

We tested whether there were differences in gene expression between species and among populations by comparing gene expression levels between *D. persimilis* and all *D. pseudoobscura* populations, and by comparing the sympatric *D. pseudoobscura* populations to our two allopatric *D. pseudoobscura.* Differences between the two species explained the largest axis of gene expression variation among both males and females (Fig 3). When each *D. pseudoobscura* population is compared to *D. persimilis* we found that all four populations had large numbers of DE genes, but gene expression differences were not consistently greater between *D. persimilis* and the sympatric populations of *D. pseudoobscura* (Supplemental Table 2. Interestingly, the *D. pseudoobscura* sympatric populations did not cluster together into a distinct group for either females or males, unlike the clustering observed between the two allopatric populations (Fig 3).

**Figure 3.**
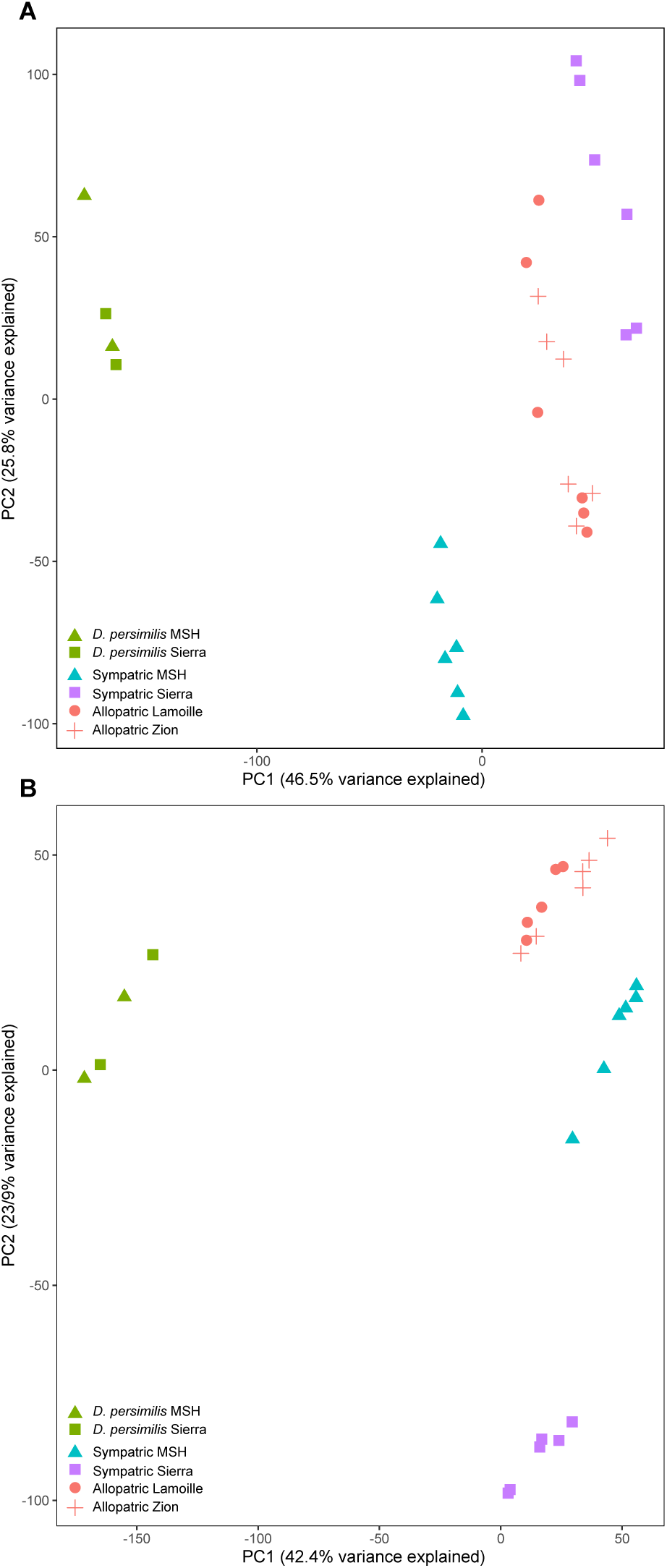
Expression differences between populations of *Drosophila pseudoobscura* and *D. persimilis* highlight that sympatric populations do not show parallel expression changes. Each point represents an isofemale line for A) adult female transcriptome and B) adult male transcriptome. The percent variance explained for each principal component is provided on the respective axis.

Each sympatric population also had numerous differentially expressed genes when contrasted with the allopatric populations. For females, the Mt. St. Helena population had 18 DE loci compared to allopatric populations while the Sierra population had 85 DE loci (Supplemental Fig 2). Of the DE expressed genes in sympatric females, only four were shared between the two comparisons; two of these genes showed higher expression and two showed lower expression in both sympatric populations (Supplemental Table 3. Four genes in common is significantly greater than the random expectation (*P=*0.025), but this is largely the result of the small number of significantly DE genes in the Mt. St. Helena population. For male expression, in individual comparisons with the allopatric populations we found 76 DE loci in the Mt St Helena population and 88 DE loci in the Sierra population (Supplemental Fig 1). Only four male loci were differentially expressed in both comparisons, which is not significantly different than the random expectation (*P*=0.56). These shared genes were differentially expressed in the same direction compared to allopatry; three genes were higher and one was lower in sympatry (Supplemental Table 4. In general, for both female and male expressed genes, the number of shared genes was only a small proportion of all DE genes identified. This indicates that the two sympatric populations have different expression profiles when compared to allopatric populations and (with few exceptions) to each other, consistent with our observation of their overall transcriptome-wide patterns of expression variation (Fig 3).

We also found that genes that were consistently DE between the two species were enriched for reproductive genes. Female genes that were DE in all four tests of *D. pseudoobscura* vs *D. persimilis* were enriched for ovary biased genes *(P<*0.001; 33 of 413 genes). Male genes that were DE in all four *D. pseudoobscura* vs *D. persimilis* comparisons, were enriched for testes biased genes (*P*<0.001; 96 of 302 genes). Reproductive gene enrichment was more heterogenous for intraspecific contrasts between each sympatric population and the allopatric populations (Supplemental Table 5), but was detected for testis-biased expression (in males; Binomial Test H0:prob>0.1627, *P*<0.0001) in the Mt. St. Helena contrast, including the known sperm competition gene *Acp53Ea*. The Sierra contrast identified enrichment in ovary-biased genes (in females; Binomial Test H0:prob>0.0165, *P*=0.0137) and also identified important spermatogenesis genes such as *sneaky* (*snky*).

### Sympatric populations show unique changes in allele frequency of reproductive genes

In general we observed that the genome wide average *Fst* was low in all pairwise *Fst* comparisons within *D. pseudoobscura* (Supplemental Table 6), which is consistent with previous estimates indicating there is little population structure across the geographical range of *D. pseudoobscura* (Fuller et al., 2016; Schaeffer & Miller, 1992). Nonetheless, there were sites within the genome that showed elevated *Fst* and subsequently elevated *PBE*. We constructed three tests to understand how these differences were distributed between *D. pseudoobscura* populations (Fig 1). Of the SNPs that were identified in both sympatric contrasts, 118 were shared PBE loci (total outliers for MSH=1801 and total outliers for Sierra=1742). The number of shared SNPs is significantly greater than the number expected at random (17 SNPs; *P*<0.001). While several of these shared SNPs had the largest PBE values (Fig 4.) these shared SNPs still represent the minority of loci that were identified in the PBE tests.

**Figure 4.**
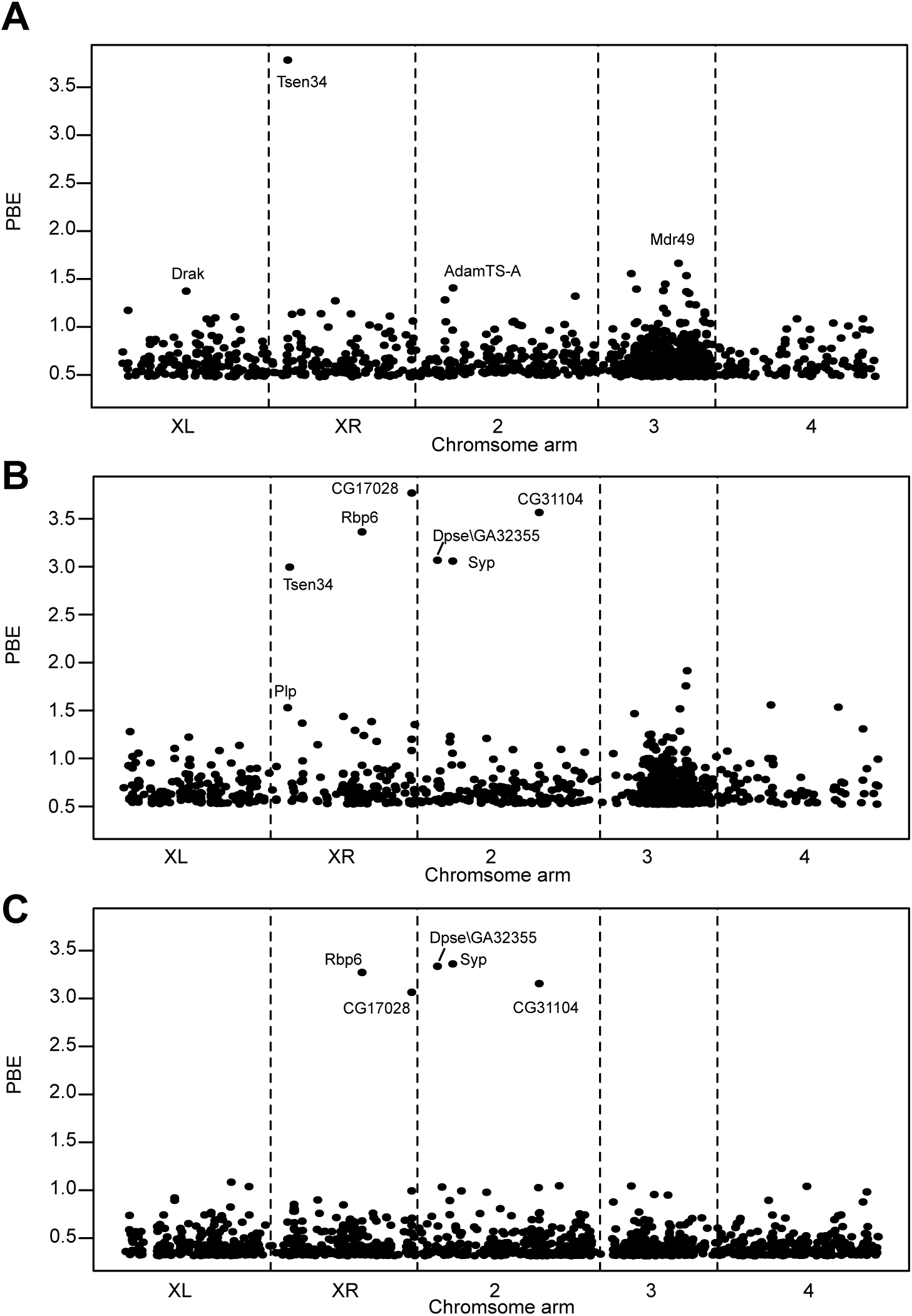
PBE outlier loci indicating SNPs under selection in sympatric populations. A) SNPs that show a signal comparing both sympatric populations to the allopatric populations. B) the Sierra sympatric population and C) the Mt St Helena sympatric population across all major chromosome arms. The labeled genes are the highest PBE outliers for each population. The name is the *D. melanogaster* ortholog, with the exception of Dpse\GA32355 that had no identifiable ortholog in *D. melanogaster*.

We tested these PBE loci for enrichment in specific biological terms and for being overrepresented in reproductive genes. In our GO analysis we found significant enrichment for nervous system development in all three tests (Supplemental Tables 7-9). Additionally for all three tests we observed strong signals of differentiation for reproductive genes (Supplemental Table 10). For the branch combining the sympatric populations, there was significant enrichment for testes biased genes (*P*=0.032) and ovary biased genes (P<0.0001). This was similar in our two tests of individual sympatric populations, where genes involved in gametogenesis were overrepresented for both the Sierra population (P<0.0001) and MSH population (P=0.00017). We also identified specific sperm competition genes within these outlier loci. Among the differences between allopatric and both sympatric populations, the genes *Est-6*, *polyph*, and *CG17575* were all outliers. For the MSH population specifically, we identified *antr, Ebp, lectin-46cb,* and *CG17575*. The gene *retn,* that is involved in female remating rate, was an outlier for each of the MSH and Sierra populations.

Lastly, to understand if any of these male reproductive genes experienced long term selection, we compared their divergence with the closely related species *D. persimilis.* We used the subset of PBE loci that had male reproductive tissue biased expression and/or were found in the proteome (n= 341 genes) and evaluated evidence of selection in these loci using MK tests. There were seven genes that showed evidence for selection after correcting for multiple testing (Supplemental Table 11). These genes were not shared between the sympatric populations. Instead different categories of genes were represented in these two populations: Genes that had male reproductive tract biased expression showed signs of positive selection in the Sierra population, whereas genes that were present in the accessory gland proteome showed signs of positive selection for the Mt. St. Helena population (Supplemental Table 11).

### Sperm competition genes are highly variable between populations

Finally, we looked at *a priori* candidate loci known to be involved in sperm competition in *D. melanogaster*, and evaluated their differentiation among populations using pairwise *Fst*—an approach that might identify differentiated genes that could not be detected as PBE outliers (see Methods). Interestingly almost all sperm competition loci had at least one SNP as an Fst outlier; some loci had outlier SNPs in every pairwise comparison (Supplemental Table 12. Compared to a random sample of the same number of loci (n=18), the sperm competition loci had an average Fst that was higher than the most differentiated SNPs in the random sample. One such gene, *aquarius* (*aqrs*), had significantly higher Fst in all pairwise tests; *aqrs* exhibited significant differences between sympatric and allopatric populations, as well as differences between sympatric populations (Supplemental Table 12). This universal differentiation among all populations would preclude identifying this gene in our previous PBE tests. Conversely, the *a priori* candidate genes that were identified as PBE outliers had a predictable pattern in pairwise Fst. For example, the gene *antares* (*antr*) was a significant outlier for PBE in the Mt. St. Helena test. There were corresponding significant Fst tests between Mt St Helena and each allopatric population, but not significant differences between allopatric populations. In contrast, *antr* was not a PBE outlier in the Sierra population contrast; as expected it also showed no significant pairwise Fst between Sierra and either allopatric population (Supplemental Table 12).

## Discussion

Reinforcement can be a strong evolutionary force, driving divergence between populations. However, for the resulting patterns of trait divergence to be stable over long periods, selection must act in specific directions to limit gene flow (Calabrese & Pfennig, 2020; Comeault & Matute, 2016; Kulmuni et al., 2020). Here we found evidence that sperm competition itself could be instrumental in maintaining reproductive differentiation among allopatric and sympatric populations of *D. pseudoobscura*, by comparing the ability of sympatric and allopatric populations to compete as simulated migrants. We also identified loci with differences in gene expression or allele frequency between sympatric and allopatric populations, including for reproductive-specific classes of genes. We found that these are most often unique to individual sympatric contexts, and include several loci known to play important roles in sperm competitive phenotypes.

Our most striking finding is that sympatric male migrants have reduced sperm competitive ability in allopatric contexts, providing a concrete mechanism that could maintain reproductive differences between allopatry and sympatry. Previously we showed that sympatric males are better than allopatric males at outcompeting heterospecifics through elevated CSP—a signal of reinforcing selection on postmating prezygotic reproductive isolation (Castillo and Moyle 2019). However, unless alleles favored in sympatry are selected against in allopatric populations, migration of these alleles should reduce and eventually eliminate this canonical sympatric versus allopatric signal of reinforcement (Calabrese & Pfennig, 2020; Comeault & Matute, 2016; Kulmuni et al., 2020). Indeed, our population genetic data confirmed that migration could in principle erase sympatric-allopatric trait differences; we observed low transcriptome-wide Fst across all our populations, consistent with the high intraspecific gene flow previously reported for *D. pseudoobscura* (Fuller et al., 2016). Nonetheless, using sperm competition phenotypes within *Drosophila pseudoobscura,* we showed that sympatric migrant males are poorer sperm competitors in allopatric contexts, providing evidence for selection specifically against these alleles in allopatry. Conversely, because sympatric males are better at outcompeting heterospecifics (Castillo and Moyle 2019), local selection for CSP in sympatry should inhibit the introduction of allopatric male genotypes into sympatric populations. In combination, these locally adapted sperm competition phenotypes provide the conditions required to create a stable pattern of reinforcement and differentiation between populations.

We expected that strong CSP and resulting interspecific sperm competition would result in divergence among sympatric and allopatric contexts in the genes underlying these traits. Indeed we observed patterns of differentiation between several specific genes implicated in sperm competition including *Est-6*, *antr*, and *CG17575*. Other detected differences also point towards roles in sperm competitive interactions. For instance, one of the extreme PBE outliers is the gene *Syp* which is a highly pleiotropic locus with roles in neural development and high expression in the testes and accessory gland (Flybase release FB2025_05; (Öztürk-Çolak et al., 2024). Moreover both genes with consistent differential expression between populations and the loci identified under selection by PBE were both enriched for reproductive categories. The strength of CSP has recently been correlated with expression differences in sperm competition genes between *Drosophila* species (Flacchi et al., 2024), indicating that expression divergence as well as protein sequence divergence could to contribute to CSP and, potentially, to sperm competition (Patlar et al., 2023). More work will need to be done on these genetic variants to determine if they play a role in sperm competition and reproductive isolation. This is especially important because we know male reproductive genes are often quickly evolving and some are *de novo* within specific lineages (Peng & Zhao, 2024; Zhao et al., 2014). Indeed, three of the genes with the largest PBE values were loci that lacked orthologs in *D. melanogaster* or had orthologs that lacked annotated biological functions.

Intriguingly, among the genetic differences between sympatry and allopatry, our analysis also identified loci enriched for behavior and neuronal development. Recent evidence strongly suggests that there are large neurological roles in females for controlling sperm competition (Chen et al., 2019; White et al., 2021), and it is possible that the enrichment for neuroanatomy and behavioral genes reflects this. One of the extreme PBE outliers was the gene *Rbp6* that is expressed in heads, and plays roles in neuronal development and in behavior (Rounds et al., 2022). Another outlier—*retained* (*retn*)—has known roles in female mating receptivity, which can contribute to sperm competition through remating rate (Laturney et al., 2018). Coincidentally *retn* activates the gene *prospero* (*pros*) (Shandala et al., 2003), which was another PBE outlier and is involved in sensory system development. Two neuronal development gene outliers, *alan shepard* (*shep*) and *Neurolgian* (*Nrg*), are also known female behavioral genes that are involved in mating discrimination (Roy and Castillo 2024). While these loci might reflect differences in other reproductive traits, they could also provide clues about mechanisms of female-side contributions to sperm competition interactions.

Over and above these specific differentiated genes, a general finding of our analyses was that each sympatric population displayed largely unique genetic differences when compared to both *D. persimilis* and to their allopatric conspecifics. Because sperm competition is dependent on the phenotypes of male competitors and females (Castillo & Moyle, 2019; Lupold et al., 2020; Tourmente et al., 2019), this creates the potential for local competitive ‘environments’ to be vastly different and dynamic, in turn generating the opportunity for local reproductive adaptation and consequent differentiation between populations (Kustra & Alonzo, 2023; Mahdjoub et al., 2023). Several pieces of evidence in our study support this broad scenario. First, sympatric males from different populations exhibited variation in their competitive phenotypes when competing with allopatric males. Second, the majority of differentially expressed genes and loci under selection were unique to individual sympatric populations rather than shared. Lastly, we detected multiple significant Fst differences in known sperm competition genes between our sympatric populations. The tendency for sperm competition to depend on specific, often idiosyncratic, male-female responses (Lupold et al., 2020; Tourmente et al., 2019) therefore appears to have generated localized patterns of evolution in our two sympatric populations, even under similar selective pressures generated by contact with *D. persimilis*.

Finally, we note that the general dependence of post-mating pre-zygotic (PMPZ) traits—like sperm competition—on local male-female co-evolution might make PMPZ differences especially likely to arise in response to reinforcing selection, and to persist after reinforcing selection has acted. If reinforcing selection acts on a single sex-specific trait, as often observed in premating isolation, this trait is able to spread across a species range following its origin in sympatry (Calabrese & Pfennig, 2020; Comeault & Matute, 2016; Kulmuni et al., 2020). However, unlike single sex-specific changes, PMPZ dynamics depend on specific interactions between male and female reproductive proteins, and both male and female components are often required for PMPZ isolation (Lupold et al., 2020; Tourmente et al., 2019). In this way, PMPZ traits might be particularly powerful mechanisms for sustaining and even amplifying reproductive differentiation between conspecific sympatric and allopatric populations, after reinforcing selection has acted.

## Notes

### Competing Interest Statement

The authors have declared no competing interest.

### Funding

- National Science Foundation: 1500911

