## Supplemental Tables and Figures for "Reinforcement and selection against migrants maintain sperm competitive and genetic differences among populations of *Drosophila pseudoobscura*"

**Supplemental Table 1.** Short Read Archive (SRA) and Drosophila Species Stock number for all isofemale strains used in this study. Libraries were sequenced across two separate runs giving two SRA numbers per strain-sex combination.

| Genotype | Male SRA Experiment numbers | Female SRA experiment numbers | Drosophila Species Stock center number |
|---|---|---|---|
| CR1 (*D. persimilis*) | SRX4676044<br>SRX4681054 | SRX4676045<br>SRX4681055 | 14011-0111.62 |
| FH2 (*D. persimilis)* | SRX4676042<br>SRX4681056 | SRX4676043<br>SRX4681057 | 14011-0111.61 |
| MSH10 (*D. persimilis)* | SRX4680973<br>SRX4680972 | SRX4680986<br>SRX4680987 | NA |
| MSH3 (*D. persimilis*) | SRX4680984<br>SRX4680985 | SRX4680990<br>SRX4680991 | 14011-0111.63 |
| MV3 (Sierra) | SRX4681027<br>SRX4681026 | SRX4681025<br>SRX4681024 | 14011-0121.276 |
| MVF18 | SRX4681023<br>SRX4681022 | SRX4681021<br>SRX4681020 | 14011-0121.277 |
| MVMS1-1 | SRX4681019<br>SRX4681018 | SRX4681030<br>SRX4681031 | 14011-0121.278 |
| MVMS1-5 | SRX4681032<br>SRX4681033 | SRX4681034<br>SRX4681035 | 14011-0121.279 |
| S4 | SRX4677328<br>SRX4681036 | SRX4677329<br>SRX4681037 | 14011-0121.274 |
| S6 | SRX4681028<br>SRX4681029 | SRX4681043<br>SRX4681042 | 14011-0121.275 |
| MSH1 (Mt St Helen) | SRX4680988<br>SRX4680989 | SRX4680982<br>SRX4680983 | 14011-0121.286 |
| MSH 4 | SRX4680993<br>SRX4680992 | SRX4680995<br>SRX4680994 | 14011-0121.287 |
| MSH 68 | SRX4680997<br>SRX4680996 | SRX4680999<br>SRX4680998 | 14011-0121.288 |
| MSH 77 | SRX4681001<br>SRX4681000 | SRX4681010<br>SRX4681011 | 14011-0121.289 |
| MSH83 | SRX4681008<br>SRX4681009 | SRX4681014<br>SRX4681015 | 14011-0121.290 |
| MSH8 | SRX4681012<br>SRX4681013 | SRX4681016<br>SRX4681017 | 14011-0121.291 |
| LC5 | SRX4681050<br>SRX4681051 | SRX4681052<br>SRX4681053 | 14011-0121.280 |

| | | | |
|---|---|---|---|
| LC8 | SRX4681048<br>SRX4681049 | SRX4680970<br>SRX4680971 | 14011-0121.281 |
| LCMS1-2 | SRX4680968<br>SRX4680969 | SRX4680966<br>SRX4680967 | 14011-0121.282 |
| LCMS2-5 | SRX4680964<br>SRX4680965 | SRX4680962<br>SRX4680963 | 14011-0121.283 |
| LCMS4-1 | SRX4680977<br>SRX4680976 | SRX4680975<br>SRX4680974 | 14011-0121.284 |
| LCMS7-3 | SRX4680981<br>SRX4680980 | SRX4680979<br>SRX4680978 | 14011-0121.285 |
| Z11 | SRX4681045<br>SRX4681044 | SRX4681039<br>SRX4681038 | 14011-0121.292 |
| Z20 | SRX4681041<br>SRX4681040 | SRX4681047<br>SRX4681046 | 14011-0121.293 |
| Z24 | SRX4680958<br>SRX4680959 | SRX4680956<br>SRX4680957 | 14011-0121.294 |
| Z36 | SRX4680954<br>SRX4680955 | SRX4680952<br>SRX4680953 | 14011-0121.295 |
| Z3 | SRX4680960<br>SRX4680961 | SRX4681007<br>SRX4681006 | NA |
| Z9 | SRX4681005<br>SRX4681004 | SRX4681003<br>SRX4681002 | 14011-0121.296 |

**Supplemental Table 2**. The number of differentially expressed genes (DE) comparing each *D. pseudoobscura* population with *D. persimilis*.

| Population (geography) | DE loci male samples | DE loci female samples |
|---|---|---|
| Zion (allopatric) | 1960 | 1135 |
| Lamoille (allopatric) | 1261 | 1202 |
| Mt St Helena (sympatric) | 471 | 1703 |
| Sierra (sympatric) | 2227 | 1030 |

**Supplemental Table 3.** The differentially expressed genes in female sympatric genotypes that were shared between both Mt. St. Helena (MSH) and Sierra populations. The Expression change in log fold change compared to allopatric populations, with the false discovery rate (FDR) for each gene.

| Gene | Expression change: MSH | FDR | Expression change: Sierra | FDR |
|---|---|---|---|---|
| FBgn0246214 | -2.4066 | 5.281e-05 | -1.8186 | 4.820e-03 |
| FBgn0080996 | 2.0617 | 5.281e-05 | 2.2174 | 8.343e-10 |
| FBgn0245828 | -2.5312 | 6.945e-04 | -1.5288 | 6.997e-02 |
| FBgn0075892 | 1.8829 | 7.602e-02 | 3.2217 | 9.348e-08 |

**Supplemental Table 4.** The differentially expressed genes in male sympatric genotypes that were shared between both Mt. St. Helena (MSH) and Sierra populations. The Expression change in log fold change compared to allopatric populations, with the false discovery rate (FDR) for each gene.

| Gene | Expression change: MSH | FDR | Expression change: Sierra | FDR |
|---|---|---|---|---|
| FBgn0072153 | 1.0441 | 0.0088 | 0.8222 | 8.537e-02 |
| FBgn0075892 | 1.5845 | 0.0469 | 2.6328 | 2.899e-09 |
| FBgn0078803 | -1.3916 | 0.0834 | -1.3630 | 6.813e-02 |
| FBgn0080996 | 1.0464 | 0.0920 | 1.3783 | 2.195e-09 |

**Supplemental Table 5**. Category enrichment for differentially expressed genes in sympatric *D. pseudoobscura* populations. The testes, RE male, ovary, and RE female are based on tissue specific expression tau statistics that we calculated (see methods). For each category we determined the proportion of genes in the genome, which was our expected proportion, and used this to compare to the proportion observed in our outlier analysis. The number of genes in each category is given in parentheses. For the MSH population there were 76 DE male genes and 29 DE female genes. For the Sierra population there were 88 DE male genes and 85 DE female genes.

| Population | Category | Expected | Observed | P-value |
|---|---|---|---|---|
| MSH | **Testes** | **16.27%** | **35.52% (27)** | **5.8e-05** |
| | RE Male | 3.08% | 3.94% | 0.51 |
| | Ovary | 1.65% | 3.44% (1) | 0.3841 |
| | RE Female | 0.34% | 0.00% (0) | 1.00 |
| Sierra | Testes | 16.27% | 18.18% (16) | 0.6644 |
| | RE Male | 3.08% | 2.22% (2) | 1.000 |
| | **Ovary** | **1.65%** | **5.88% (5)** | **0.0137** |
| | RE Female | 0.34% | 0.00% (0) | 1.00 |

**Supplemental Table 6.** Genome average pairwise Fst between *D. pseudoobscura* populations demonstrating high levels of gene flow across the species range.

| | Zion | Lamoille | Mt St Helena | Sierra |
|---|---|---|---|---|
| Zion | | | | |
| Lamoille | 0.066 | | | |
| Mt St Helena | 0.027 | 0.075 | | |
| Sierra | 0.037 | 0.083 | 0.0283 | |

**Supplemental Table 7.** The gene ontology (GO) terms that were overrepresented for the test that identified loci under selection common to the sympatric populations, using the PBE statistic. The gene names are the *D. melanogaster* orthologs for the *D. pseudoobscura* genes. These categories were significant after testing for multiple correction.

| GO term | Number of genes | P-value | Genes |
|---|---|---|---|
| lipid transport | 15 | 1.1701E-05 | Itpr,GIIIspla2,CG4301,CG33298,CG9084,ATP8B,Nrx-1,lilli,scramb1,CG34120,apolpp,CG42237,ATP8A,CG42668,Fatp2 |
| cell surface receptor signaling pathway | 66 | 4.1494E-05 | arm,arr,Dab,dpp,E(bx),fz,Galphao,gro,Pp1alpha-96A,shn,sl,Egfr,ttk,RYa-R,babo,fra,CkIalpha,Diap2,apt,stet,stumps,Dad,Pp4-19C,CG8910,nito,PPP4R2r,Socs16D,Pvf1,CG14234,cactin,rau,Uvrag,Myd88,Mmp2,wdp,Xxylt,MAN1,rno,Hipk,Herc4,Fit1,spartin,Tsp86D,Sdr,puf,CG11318,kek6,Trim9,Ddr,akirin,apolpp,qvr,dl,Ubr3,clos,hpo,nej,cic,pyd,hppy,Uba3,sli,rdx,sno,Doa,lola |
| nervous system development | 103 | 4.504E-05 | arm,arr,ash2,brm,ck,Dab,dpp,dsx,EcR,fz,DCTN1-p150,Galphao,gro,Poxm,shg,shn,sl,Egfr,ttk,betaTub60D,Gad1,fs(1)h,Itpr,hig,Spn,babo,fra,insc,shot,Nrx-IV,cher,Pi3K59F,SMC3,apt,ast,stumps,dom,Dad,NiPp1,drosha,Amph,Smyd4-4,ced-6,lva,rau,Uvrag,Mcm10,Smyd4-1,Mmp2,dila,tou,sug,CG6701,Ntan1,Efhc1.2,Mgat1,wdp,Liprin-gamma,PIP5K59B,IntS1,Mmp1,rno,Ctr9,spartin,Tsp86D,Arfip,Lrrk,Nrx-1,SMC1,robo3,lilli,olf186-F,bchs,fdl,nolo,Trim9,shep,Mical,spri,mute,pea,dl,par-1,stj,hpo,pdm3,nej,Dhc64C,aPKC,AGO1,Patronin,Dark,sli,how,CG44153,PIG-V,sno,Doa,unc-104,PI4KIIIalpha,Pak,wupA,lola |
| intracellular signal transduction | 59 | 0.0002301 | dpp,Galphao,sl,Egfr,Rab3,trpl,Spn,PhKgamma,Dp,Pi3K59F,Pkn,Mekk1,Rep,Wdr24,Socs16D,Pvf1,RhoGAP19D,Myd88,CG5522,swi2,Dlish,CG10737,sigmar,MAN1,Hipk,Herc4,anchor,Gyc88E,Sur-8,Tusp,Capa,CG11318,CG17698,ACXE,olf186-F,GEFmeso,Unc-89,CG34384,spri,CG42260,Ziz,dl,par-1,Opa1,grp,hpo,Exn,rdgA,aPKC,CG43102,hppy,Stacl,spoon,rdx,rdgC,Doa,Pde8,PI4KIIIalpha,Pak |
| response to external stimulus | 89 | 0.0004369 | Arr1,brm,car,Dab,dpp,dsx,E(bx),EcR,fz,DCTN1-p150,Galphao,lt,Pp1alpha-96A,shg,betaTub60D,nonA,Gad1,trpl,Itpr,babo,fra,Mef2,insc,mtd,shot,Calx,cher,tim,Diap2,Pi3K59F,ktub,dom,Dad,btv,ssx,CBP,NiPp1,Wdr24,ECSIT,Myd88,Mmp2,polyph,sug,Pepck2,PIP5K59B,CG9380,Hipk,Herc4,CPT2,Fit1,Dnah3,Adgf-A,Sdr,Lrrk,puf,brv3,CG13551,wun2,robo3,TotX,Trim9,shep,Sec16,Mical,pain,EMRE,akirin,shakB,Camta,dl,stj,Exn,rdgA,pdm3,Dhc64C,LanB1,aPKC,Ect4,Dark,sli,CG44153,Trpm,rdgC,Doa,unc-104,Pak,PPO1,Phm,lola |

**Supplemental Table 8.** The gene ontology (GO) terms that were overrepresented for the test that identified loci under selection specific to the MSH sympatric population, using the PBE statistic. The gene names are the *D. melanogaster* orthologs for the *D. pseudoobscura* genes. These categories were significant after testing for multiple correction.

| GO term | Number of genes | P-value | Genes |
|---|---|---|---|
| cell motility | 40 | 4.344E-05 | dpp,EcR,ena,tor,ttk,18w,faf,fra,CASK,tilB,Rme-8,Vang,apt,RN-tre,Sin3A,Fpps,Ptpmeg2,CG10958,CG11178,CG1486,mip120,CG6805,Lrt,wun2,tai,Vhl,slam,Plp,kuz,par-1,kug,Dhc64C,Msp300,Hmgcr,dally,how,Src42A,PDZ-GEF,Fhos,Sec6 |
| intracellular signal transduction | 58 | 7.8874E-05 | ca,dpp,ImpL2,Rab32,sev,tor,Ac76E,Rab3,mof,Pdk1,Pkn,Uba1,Sbf,Rgl,CG18659,Smurf,Rab3-GEF,Socs16D,RhoGAP19D,Myd88,egr,Mos,RhoGAP54D,P32,swi2,CG10737,Sesn,MAN1,Usp5,RhoGDI,Gyc88E,PKD,Tusp,CycG,CG17698,Aplip1,GEFmeso,RabX4,CG33958,Dgk,Lgr4,tum,C3G,sqa,gwl,par-1,raskol,S6kII,hppy,Lpin,Hmgcr,Stacl,Lst8,Src42A,PDZ-GEF,Doa,Pde8,wap |
| nervous system development | 93 | 0.0004349 | arr,brm,Dab,dpp,dsx,EcR,ena,ImpL2,klar,mle,Poxm,sev,shn,sm,ttk,yrt,uzip,18w,Fur1,retn,noc,Itpr,hig,brat,dap,cora,kermit,babo,fra,insc,CASK,cnn,chn,Vang,apt,loco,dom,Pdk1,Uba1,Amph,Smyd4-4,lva,Tbce,p47,egr,CG12935,Smyd4-3,tou,Efhc1.2,Mgat1,Liprin-gamma,enok,Mmp1,Eps-15,rno,Usp5,MED10,Mau2,RhoGAP92B,Lrrk,Syp,Dcr-1,Prosap,tai,dpr11,nub,mute,Snoo,tum,beta-Spec,cwo,kuz,gkt,par-1,Vmat,stj,didum,pdm3,Dhc64C,CG42797,nrm,AGO1,S6kII,Acsl,scrib,dally,how,Src42A,Nrg,PDZ-GEF,Doa,Sec6,lola |
| cell morphogenesis | 53 | 0.0005169 | arr,brm,Dab,dpp,dsx,EcR,ena,ImpL2,mle,Poxm,shn,sm,ttk,uzip,retn,brat,kermit,babo,fra,cnn,chn,Vang,colt,dom,Ankle2,lva,Liprin-gamma,enok,Slik,Mau2,Lrrk,Dcr-1,CG41099,Prosap,tai,dpr11,nub,Snoo,tum,beta-Spec,cwo,kuz,pdm3,Dhc64C,CG42797,Acsl,scrib,dally,Src42A,Nrg,PDZ-GEF,Fhos,lola |
| epithelium development | 85 | 0.0005175 | arr,brm,salr,Dab,dpp,dsx,EcR,ena,ix,lin,mus304,shn,Su(var)2-10,tor,ttk,yrt,kst,18w,fs(1)N,noc,dap,cora,kermit,babo,Mef2,insc,CASK,Rme-8,Vang,apt,vkg,colt,loco,dom,RN-tre,Pkn,Sin3A,Uba1,SPARC,Ptpmeg2,Set2,RhoGAP19D,Irk3,mip120,CG8405,RhoGAP54D,Slik,Mmp1,CG11399,RhoGAP92B,Syp,Mettl3,CycG,CG41099,tai,Vhl,CG30069,GEFmeso,fred,CG31926,Drak,Idgf5,psd,sqa,kuz,par-1,clos,kug,Dhc64C,Fs(2)Ket,S6kII,scrib,Lpin,Hmgcr,dally,Ack-like,how,Src42A,Nrg,PDZ-GEF,Fhos,Sec6,Phm,Pal1,lola |

**Supplemental Table 9.** The gene ontology (GO) terms that were overrepresented for the test that identified loci under selection specific to the Sierra sympatric population, using the PBE statistic. The gene names are the *D. melanogaster* orthologs for the *D. pseudoobscura* genes. These categories were significant after testing for multiple correction.

| GO term | Number of genes | P-value | Genes |
|---|---|---|---|
| nervous system development | 123 | 4.80006398996627e-8 | asp,boss,brm,DCTN1-p150,Got2,gro,klar,E(spl)mbeta-HLH,mr,sesB,svp,tkv,Tm1,Egfr,trx,ttk,Ptp4E,Fur1,Syn,pros,zfh2,fs(1)h,retn,tok,tws,gig,Mbs,robo1,Itpr,brat,Akt1,Spn,Moe,Snr1,Arp2,Hem,shot,Nrx-IV,Rel,cher,Nf1,chn,ksr,SMC3,nrv2,ast,spen,Cbl,Dredd,Su(z)12,chb,RhoGEF2,Tip60,Rok,Nipped-B,NiPp1,tacc,Rip11,jim,TBCD,drpr,Dlic,daw,rau,Pex19,CdGAPr,Tbce,Smyd4-3,Ptp52F,Liprin-gamma,CG13531,Usp5,slow,Pura,Smyd4-2,Cep135,wnd,Wnk,Syp,Usp8,Dcr-1,pasha,Gclc,Prosap,bchs,gogo,shep,dpr11,Mical,Nlg4,spri,mute,sif,Snoo,Invadolysin,mbm,psidin,beta-Spec,crb,zld,cwo,kuz,Vmat,inaE,pdm3,nej,Npc1b,Cpn,Dhc64C,sdt,CG42797,ana1,nrm,S6kII,sick,dally,Src42A,PDZ-GEF,Doa,Nsf2,kis,PI4KIIIalpha,Sac1 |
| actin cytoskeleton organization | 40 | 4.0933E-06 | bru1,asp,capu,flw,kel,Prm,spir,Tm1,fwd,svr,Gel,Spn,Moe,Arp2,Hem,shot,cher,Mlp84B,RhoGEF2,Ocrl,Rok,CG5023,CG30183,CG32264,Mical,sif,psidin,crb,scra,kug,brun,Msp300,CG43102,sick,Ack-like,Lst8,Src42A,ghi,btsz,PI4KIIIalpha |
| synapse organization | 40 | 4.8397E-06 | Got2,mr,sesB,tkv,Fur1,Syn,pros,gig,brat,Akt1,Spn,Arp2,Hem,Gem3,Nrx-IV,Nf1,RtGEF,Tip60,drpr,Upf2,Tbce,Liprin-gamma,wnd,dpr9,Syp,Prosap,dpr6,bchs,dpr11,Mical,Nlg4,beta-Spec,nej,S6kII,Src42A,Neto,Nsf2,kis,PI4KIIIalpha,Sac1 |
| epithelium development | 104 | 1.1257E-05 | brm,capu,dec-1,flw,DCTN1-p150,kel,px,spir,svp,tkv,Egfr,trx,ttk,Vm26Ab,Ptp4E,fwd,zfh2,svr,tok,tws,bowl,fs(1)M3,Mbs,robo1,Moe,Snr1,Trl,shot,Nrx-IV,Su(Tpl),cher,Cp7Fb,Mlp84B,ksr,Rme-8,SMC3,ast,lama,vkg,spen,colt,Cbl,lig,crol,RhoGEF2,mip130,bab2,Rok,Idgf4,drpr,Cdc7,CHES-1-like,xit,Set2,RhoGAP19D,daw,Irk3,mip120,slow,Pura,CG11399,Wnk,Syp,Usp8,Mettl3,CycG,pasha,Gclc,cbt,fred,Drak,Idgf5,spri,Invadolysin,psidin,sqa,crb,zld,kuz,kug,nej,Npc1b,Dhc64C,sdt,Csk,magu,S6kII,nclb,Lpin,Hmgcr,dally,Ack-like,rdx,Src42A,Irk1,cv-d,PDZ-GEF,Nsf2,kis,Sec5,btsz,Mcr,Syx7,Sac1 |
| cell differentiation | 162 | 2.5138E-05 | abo,Act5C,Mps1,bru1,asp,boss,brm,capu,dec-1,flw,DCTN1-p150,kel,klar,mei-9,E(spl)mbeta-HLH,mus312,Pfk,Prm,sesB,spir,svp,tkv,Tm1,Egfr,trx,ttk,Vm26Ab,Ptp4E,Fur1,pros,fs(1)h,retn,tok,tws,gig,fs(1)M3,Mbs,robo1,faf,Itpr,brat,Akt1,Spn,ms(2)35Ci,Moe,Snr1,twin,Hem,Trl,shot,Nrx-IV,Rel,cher,Cp7Fb,Mlp84B,Nf1,chn,ksr,Rme-8,SMC3,nrv2,ast,lama,spen,colt,Cbl,mspo,lig,Dredd,Su(z)12,chb,Zfrp8,RhoGEF2,mip130,bab2,Rok,Sara,NiPp1,tacc,Rip11,jim,TBCD,goe,drpr,Cdc7,Pat1,xit,Dlic,tho2,daw,rau,uif,CdGAPr,Smyd4-3,mip120,Ptp52F,Liprin-gamma,CG13531,Usp5,slow,Pura,Smyd4-2,Cep135,ATPsynbetaL,wnd,Wnk,aux,Syp,Usp8,Dcr-1,Mettl3,pasha,Gclc,Prosap,wun2,bchs,gogo,shep,dpr11,Mical,spri,sif,Snoo,AGO2,psidin,beta-Spec,crb,zld,out,cwo,kuz,Vmat,asl,inaE,lqfR,scra,kug,pdm3,nej,Cpn,mi,Dhc64C,Msp300,sdt,CG42797,Csk,ana1,nrm,S6kII,milt,nclb,sick,dally,Cyt-b5,Src42A,PDZ-GEF,Doa,kis,Sec5,PI4KIIIalpha,rump,Sac1 |
| cell morphogenesis | 64 | 3.5683E-05 | brm,DCTN1-p150,tkv,Tm1,trx,ttk,Ptp4E,pros,fs(1)h,retn,tok,Mbs,robo1,brat,Moe,Snr1,Arp2,Hem,shot,Rel,cher,chn,spen,colt,chb,RhoGEF2,Rok,NiPp1,jim,TBCD,qsm,xit,Dlic,daw,CdGAPr,Ptp52F,Liprin-gamma,CG13531,Pura,wnd,Wnk,Dcr-1,Gclc,Prosap,gogo,dpr11,Mical,spri,sif,Snoo,psidin,beta-Spec,crb,cwo,kuz,pdm3,Dhc64C,CG42797,sick,dally,Src42A,PDZ-GEF,kis,Sac1 |

**Supplemental Table 10.** Category enrichment for PBE outliers in sympatric *D. pseudoobscura* populations. The gametogenesis category is based on orthologous genes in *D. melanogaster* that have been identified with that GO term. The testes, RE male, ovary, and RE female are based on tissue specific expression tau statistics that we calculated (see methods). For each category we determined the proportion of genes in the genome, which was our expected proportion, and used this to compare to the proportion observed in our outlier analysis. The number of genes in each category is given in parentheses. Sympatry indicates shared SNPs that differentiated both populations from allopatric populations.

| Population | Category | Expected | Observed | P-value |
|---|---|---|---|---|
| M | **Gametogenesis** | **8.81%** | **12.74% (98)** | **0.00017** |
| | Testes | 16.27 | 14.46 (198) | 0.9698 |
| | RE Male | 3.08 | 2.77 (38) | 0.7689 |
| | **Ovary** | **1.65** | **9.42 (129)** | **<0.0001** |
| | RE Female | 0.346 | 0.584 (8) | 0.1071 |
| S | **Gametogenesis** | **8.81%** | **13.8 (123)** | **6.223e-07** |
| | Testes | 16.27 | 16.33 (219) | 0.496 |
| | RE Male | 3.08 | 2.23 (30) | 0.976 |
| | **Ovary** | **1.65** | **8.65 (116)** | **<0.00001** |
| | RE Female | 0.346 | 0.37 (5) | 0.4946 |
| Sympatry | **Gametogenesis** | **8.81%** | **11.49 (94)** | **0.00545** |
| | **Testes** | **16.27** | **18.24 (241)** | **0.0302** |
| | RE Male | 3.08 | 2.87 (38) | 0.6933 |
| | **Ovary** | **1.65** | **5.22 (69)** | **<0.00001** |
| | RE Female | 0.346 | 0.37 (5) | 0.4816 |

**Supplemental Table 11.** Summary of loci that were PBE outliers that also had signals of adaptive evolution from McDonald-Kreitman tests, comparing fixed and polymorphic substitutions with *D. persimilis* data that we had collected.

| *D. pseudoobscura* FBgn | *D. melanogaster* ortholog | Male RT tissue specific | Present in Accessory proteome | Population | P-value (FDR) |
|---|---|---|---|---|---|
| FBgn0071006 | *CG11369* | Yes | No | Sierra | <0.0001 (0.0022) |
| FBgn0244480 | *JYalpha* | Yes | Yes | Sierra | <0.0001 (0.0002) |
| FBgn0244714 | - | Yes | No | Sierra | 0.0003 (0.0079) |
| FBgn0077126 | *nonC* | No | Yes | MSH | 0.0009 (0.0305) |
| FBgn0077749 | *Marf* | No | Yes | MSH | <0.0001 (0.0036) |
| FBgn0078484 | *tyf* | No | Yes | MSH | 0.0001 (0.0059) |
| FBgn0244181 | *Ubr3* | No | Yes | MSH | <0.0001 (0.0037) |

**Supplemental Table 12**. Allele frequency differences of genes that are orthologs of known sperm competition genes in *D. melanogaster*. The reported value is the maximum *Fst* for a given locus in each pairwise comparison. Values that are bolded are *Fst* outliers, with the 95% cutoff reported at the bottom of the table. The last row of the table is the mean *Fst* for the maximum values for all sperm competition genes across a population comparison. These were all significantly greater (P<0.05) than a random sample of genes.

| *D. melanogaster* ortholog | *D. pseudoobscura* FBgn | M vs L | M vs Z | L vs Z | S vs L | S vs Z | S vs M |
|---|---|---|---|---|---|---|---|
| *Gld* | FBgn0012699 | **0.6341** | **0.6756** | 0.2333 | **0.3939** | **0.393** | **0.3939** |
| *Sems* | FBgn0070474 | 0.1870 | 0.0250 | 0.2000 | 0.3600 | **0.3200** | 0.1000 |
| *Abd-B* | FBgn0071172 | 0.2500 | 0.0570 | 0.1548 | **0.4127** | 0.2500 | **0.3478** |
| *intr* | FBgn0071743 | **0.7735** | **0.3637** | **0.3478** | **0.4333** | **0.345** | **0.7735** |
| *CG15200* | FBgn0073600 | 0.3000 | 0.0000 | 0.3000 | 0.1666 | 0.2500 | **0.3181** |
| *lectin-46Cb* | FBgn0074106 | **0.3722** | **0.6250** | **0.5764** | 0.3141 | **0.4166** | **0.5185** |
| *CG17575* | FBgn0074591 | **0.6138** | **0.7717** | 0.1000 | 0.2500 | **0.3200** | **0.3853** |
| *Ebp* | FBgn0075435 | **0.4750** | **0.6000** | **0.4000** | **0.4000** | **0.3000** | **0.4782** |
| *frma* | FBgn0076878 | **0.6367** | **0.4387** | **0.5428** | **0.5767** | 0.2830 | 0.2500 |
| *Acp62F* | FBgn0078773 | 0.2500 | 0.1765 | **0.3617** | 0.2500 | **0.4000** | **0.3600** |
| *hdly* | FBgn0079015 | **0.4666** | **0.4047** | **0.4666** | **0.5043** | **0.3727** | **0.5416** |
| *Esp* | FBgn0080019 | **0.4333** | 0.2500 | **0.3853** | 0.3600 | **0.3853** | **0.3200** |
| *Nep2* | FBgn0082000 | 0.3454 | **0.3333** | **0.5043** | **0.6500** | **0.4500** | **0.3600** |
| *CG9997* | FBgn0082155 | 0.2500 | 0.2000 | **0.5764** | **0.6250** | **0.6250** | 0.2285 |
| *Est-6* | FBgn0243549 | **0.5615** | **0.4500** | **0.4666** | **0.7111** | **0.7111** | **0.6000** |
| *Est-6* | FBgn0243571 | **0.5384** | 0.0000 | 0.3200 | NA | 0.2837 | **0.5000** |
| *Acp53Ea* | FBgn0243575 | 0.0000 | 0.1773 | **0.3992** | 0.0570 | 0.0769 | 0.0000 |
| *CG17450* | FBgn0244291 | **0.4666** | **0.3600** | **0.5428** | **0.4666** | 0.2000 | 0.2909 |
| *Pkd2* | FBgn0245539 | 0.0000 | 0.1333 | 0.1111 | 0.2027 | 0.2767 | 0.2767 |
| *antr* | FBgn0245599 | **0.4782** | **0.6250** | 0.3000 | 0.1666 | 0.2000 | **0.3000** |

| | | | | | | | |
|---|---|---|---|---|---|---|---|
| *lectin-46Ca* | FBgn0245732 | **0.4666** | **0.5714** | 0.2285 | **0.4000** | 0.3000 | **0.3000** |
| *Pkd2* | FBgn0247109 | 0.2837 | **0.3560** | 0.3200 | **0.4000** | **0.5764** | **0.4387** |
| *Pp1-13C* | FBgn0247960 | **0.4666** | 0.1483 | **0.3600** | **0.4782** | 0.2500 | **0.3853** |
| *aqrs* | FBgn0248361 | **0.5428** | 0.1333 | **0.5428** | **0.4333** | **0.3727** | **0.3560** |
| *Dnah3* | FBgn0249901 | **0.7058** | **0.4000** | **0.8000** | **0.5043** | **0.5428** | **0.4988** |
| *SP* | FBgn0270940 | **0.4750** | 0.1000 | 0.3200 | 0.1333 | **0.5714** | **0.3600** |
| 95% cutoff for each population comparison | | 0.3600 | 0.3037 | 0.3454 | 0.3727 | 0.3000 | 0.3000 |
| Average maximum Fst | | 0.4220 | 0.3221 | 0.3792 | 0.3860 | 0.3644 | 0.3724 |

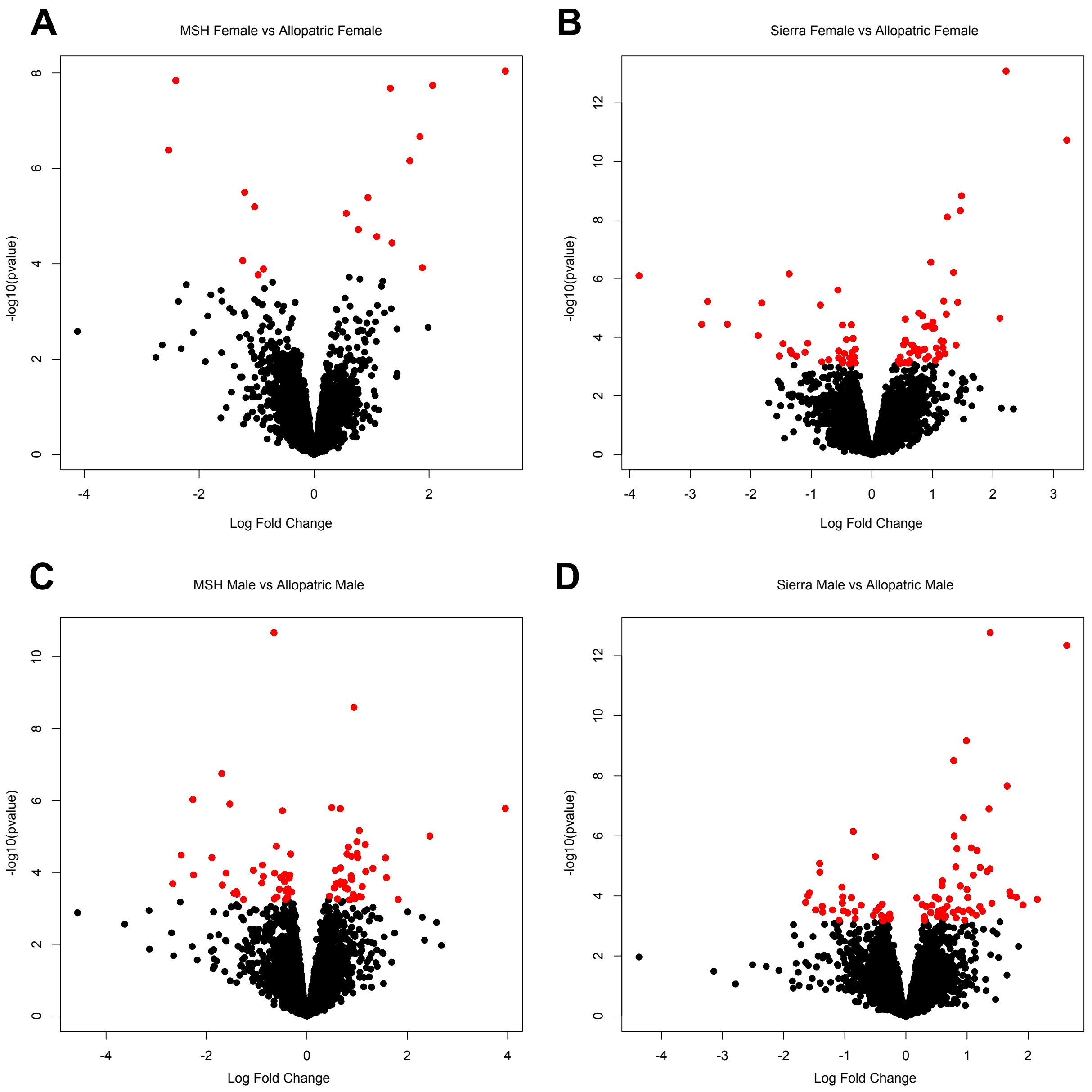


Supplemental Figure 1. Differences in gene expression between sympatric and allopatric populations of *D. pseudoobscura*. A) MSH females compared to allopatric females. B) Sierra females compared to allopatric males. C) MSH males compared to allopatric males. D) Sierra males compared to allopatric males. In each panel red dots are significant after controlling for multiple tests.
